# Structural reorganization and genomic context define a divergent lineage of the *Wolbachia* male-killing gene *wmk*

**DOI:** 10.64898/2026.03.16.712054

**Authors:** Ranjit Kumar Sahoo

## Abstract

Patterns of diversity in symbiont effector genes provide insight into the evolutionary processes that shape their diversification, particularly those arising from host–symbiont interactions. In one of the most widespread symbiont genera, *Wolbachia*, the male-killing candidate gene *wmk* encodes a putative transcriptional regulator. Sequence divergence of this effector gene from a limited number of strains has revealed at least five phylogenetic types. However, additional *wmk* variants characterized by a large inframe deletion and protein reorganization suggest that diversity in *wmk* extends beyond sequence variation alone. To gain further insight into *wmk* effector diversity, homologous proteins from 251 *Wolbachia* genomes were analyzed using comparative sequence and structure-informed approaches. The results show that sequence and structural diversification largely follow similar patterns; however, one lineage newly identified in this analysis stands out due to pronounced structural reorganization. The distinct genomic neighborhood of this divergent lineage, relative to other *wmk* lineages, suggests additional diversity at the regulatory level. Together, these findings demonstrate that variation in protein structure and genomic context complements sequence-level polymorphism in shaping *wmk* effector diversity in *Wolbachia*. Further analyses indicate that symbiont supergroup and host taxonomic order constrain the distribution of the divergent *wmk* lineage.

## Introduction

Effector proteins secreted by bacterial symbionts are key mediators of host–symbiont co-evolution (Muller et al., 2026). These proteins manipulate host cellular processes to facilitate infection, persistence, or transmission. In antagonistic contexts, effectors continually evolve to adapt to host targets and evade host defenses (Martel et al., 2021; Derbyshire and Raffaele, 2023). Such reciprocal adaptation can drive rapid diversification of effector genes, generating extensive sequence variation (Muller et al., 2026; Liu et al., 2019). Yet many effectors retain conserved structural frameworks that preserve interactions with host targets (Seong and Krasileva, 2023; Konaté et al., 2019). In some cases, however, sequence divergence can lead to structural reorganization (Williams and Lovell, 2009), potentially generating new functional properties or enabling diversification of effector activity. Identifying structurally divergent effector variants and examining their distribution across symbiont lineages and in association with host taxa offers an opportunity to understand how co-evolutionary forces shape effector diversity in intracellular symbionts.

Among intracellular bacterium, *Wolbachia* is the most widespread infecting almost half of arthropod species (Weinert et al., 2015). In these hosts, its effects often manifest as reproductive manipulations such as cytoplasmic incompatibility, male killing, feminization of genetic males, and induction of parthenogenesis (Kaur et al., 2021). These phenotypes enhance maternal transmission of the bacterium and promote its persistence within host populations, while often imposing substantial fitness costs on hosts (Sahoo 2016; Engelstädter and Hurst, 2009). Molecular studies indicate that several of these phenotypes are mediated by bacterial effectors that interact with host DNA or proteins, disrupting key developmental processes during embryogenesis (Li et al., 2024; Permutter et al., 2019; Beckmann et al. 2017; LePage et al., 2017). Consistent with adaptation to the intracellular eukaryotic environment, many *Wolbachia* effectors are enriched in eukaryotic-like domains – such as ankyrin repeats and leucine-rich motifs – which facilitate interactions with host signaling pathways (Hamilton et al., 2025; Hamilton and Newton, 2024; Rice et al., 2017; Kedzierski et al., 2004). For example, the male-killing factor Oscar in *w*Fur strain (host: *Ostrinia furnacalis*) contains numerous ankyrin repeats and targets the host masculinization factor Masc, disrupting male development and causing embryonic male lethality (Katsuma et al., 2022).

Among the male-killing factors identified in *Wolbachia*, the most widely distributed candidate is encoded by the WO-mediated killing (*wmk*) gene (Permutter et al., 2019), which is the focus of the present study. *wmk*-encoded effector is predicted to function as a transcriptional regulator and typically contains two helix–turn–helix (HTH) domains separated by an extended inter-domain region, a configuration consistent with DNA-binding proteins (Sahoo et al., 2026; Permutter et al., 2019; see also Lefoulon et al., 2025). Although the precise molecular mechanism remains unresolved, evidence from natural infections and transgenic assays suggests that Wmk interferes with the host dosage compensation machinery during male embryonic development (Permutter at al., 2019; Harumoto et al., 2018; Riparbelli et al., 2012). Comparative analyses have identified substantial sequence diversity among Wmk homologs, which cluster into five phylogenetic clades (Lefoulon et al., 2025). In addition to these canonical forms, a small number of highly divergent homologs have been identified that display extensive sequence divergence accompanied by structural remodeling (Sahoo et al., 2026). Whether these variants represent peripheral divergence within existing Wmk diversity or instead constitute a distinct evolutionary lineage remains unresolved.

To gain insight into how sequence and structural diversification shape overall Wmk effector diversity, homologous proteins from 251 *Wolbachia* genomes were examined using sequence- and structure-informed analyses alongside genomic neighborhood comparisons. The association between *Wolbachia* supergroup, host order, and the distribution of Wmk was further evaluated.

## Results and Discussions

### 2HTH-Wmk across Wolbachia

To characterize the repertoire of Wmk homologs, 251 *Wolbachia* genomes spanning nine supergroups were analyzed (Fig. 1A, B). These strains infect 224 host species distributed across 15 arthropod orders and the order Rhabditida (Nematoda) (Fig. 1B). The genome assemblies were highly complete, with BUSCO completeness scores ranging from ~80.6% to 99.7% (median= 98.8%) (Fig. 1C). Amino acid sequences of all predicted coding regions (annotated using Prokka) were screened to identify Wmk homologs across the 251 genomes using MMSeqs2. Three previously characterized Wmk variants – *w*Mel-Wmk (AAS14326), *w*Bif-Wmk (QEQ51101), and *w*Zbi-Wmk (XXB89297) – sharing 26–48% pairwise identity, were used as reference queries (Sahoo et al., 2026). After removal of redundant hits arising from multiple matches (see Methods), the final dataset retained only full-length variants containing two helix–turn–helix (HTH) domains (Fig. 1A). These full-length homologs are designated here as 2HTH-Wmk (hereafter referred to as Wmk unless otherwise specified).

**Figure 1.**
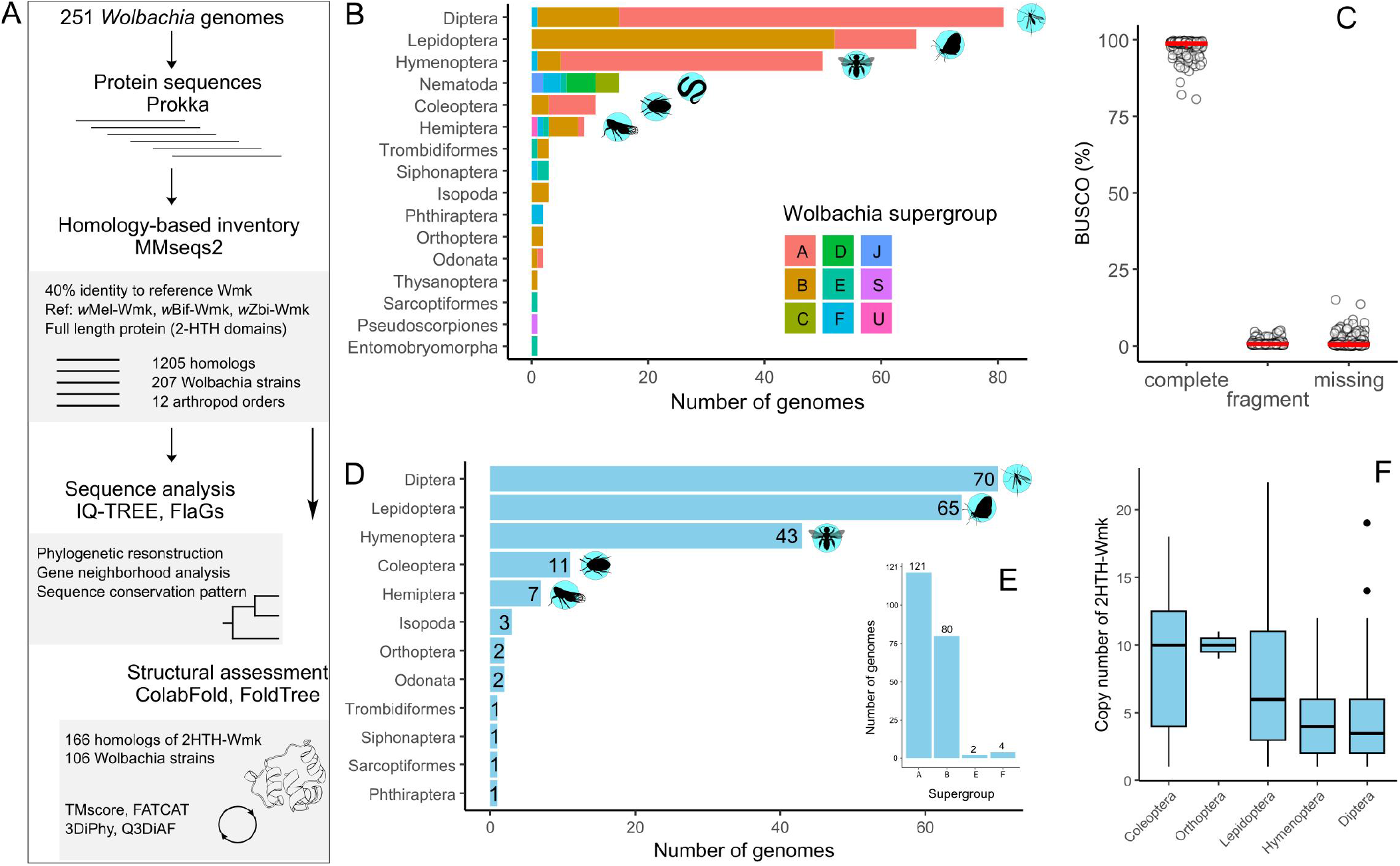
Workflow and genomic distribution of full-length Wmk homologs across *Wolbachia* genomes. (A) Schematic representation of the analytical workflow. Annotated protein sequences from 251 *Wolbachia* genomes were screened for full-length Wmk homologs using MMSeqs2. Retrieved homologs were subjected to phylogenetic reconstruction, residue conservation analysis, and genomic neighborhood mapping. Predicted structures for a representative subset were further compared to evaluate phylogenetic concordance and global structural divergence. (B) Bar plot showing supergroup-wise distribution of *Wolbachia* strains across host orders. (C) Genome completeness assessment of the 251 *Wolbachia* assemblies using BUSCO against the Rickettsiales_odb10 dataset. Each circle represents a single genome. Red box plots denote the median and interquartile range. (D) Host order-wise distribution of 207 *Wolbachia* genomes identified to encode full-length Wmk homologs. Numbers above bars indicate genome counts. (E) Supergroup-wise distribution of the same 207 genomes harboring full-length Wmk homologs, with bar labels indicating genome counts. (F) Distribution of per-genome copy number of full-length Wmk homologs. Genomes are grouped by host order; the five most represented host orders are shown.

To evaluate the effectiveness of the bioinformatics pipeline in detecting 2HTH-Wmk homologs, the repertoire identified in 30 selected *Wolbachia* genomes was compared with that reported by Lefoulon et al. (2025), in which these genomes were deeply curated. The number of 2HTH-Wmk homologs matched exactly in 16 genomes (~53.3%) (Table S1). In the remaining 14 genomes, the count was consistently lower by one homolog relative to Lefoulon et al. (2025). These discrepancies likely reflect the stricter filtering criteria applied in the present analysis, which retained only intact loci containing complete HTH domains to minimize false-positive identification. Homologs bearing pseudogenized regions or only partially overlapping one or both HTH domains relative to the reference sequences were excluded. For example, the previously described truncated Wmk locus (HCR15_02295) in *w*Bif (Lefoulon et al., 2025) was not recovered in the current dataset. Similarly, one locus each in *w*Sca and *w*Mau was excluded due to incomplete overlap with the reference HTH domains.

Analysis of the Wmk repertoire across 251 genomes revealed that ~83% of *Wolbachia* strains (207/251) harbor 2HTH-Wmk homologs (Fig. 1D). When adjusted for potential Type II error arising from incomplete genome assemblies (based on assembly type “linear” or contig number >1), prevalence increased to ~91% (207/227). The 207 Wmk-positive strains predominantly belong to supergroups A and B (~97%) and infect hosts spanning 12 arthropod orders (Fig. 1D, 1E). Notably, 2HTH-Wmk was not detected in strains infecting filarial nematodes (order Rhabditida; n= 15), consistent with their obligate mutualistic association with these hosts (Taylor et al., 2013). Exclusion of these mutualistic strains further elevated estimated prevalence to ~94% (207/220). This dataset represents the first large-scale quantitative assessment of Wmk distribution across *Wolbachia* genomes, providing empirical support for prior predictions of its widespread occurrence (Perlmutter et al., 2019). Consistent with the present analysis, a previous survey of 32 *Wolbachia* genomes from four arthropod host orders reported ~94% prevalence of 2HTH-Wmk (Lefoulon et al., 2025). Taken together, these findings suggest that Wmk, in its broader variant spectrum – including single HTH-containing forms (1HTH-Wmk) – may be present in the majority of *Wolbachia* genomes.

In addition to filarial nematodes, 2HTH-Wmk was not detected in strains infecting three arthropod orders: Thysanoptera (thrips; n= 1), Entomobryomorpha (springtails; n= 1), and Pseudoscorpiones (n= 1). However, absence in these orders remains inconclusive due to their limited sampling in the current dataset (a single representative species per order) and, in the case of Pseudoscorpiones, incomplete genome assembly. Among the 207 *Wolbachia* strains harboring 2HTH-Wmk, ~86% infect only three arthropod orders: Diptera (flies; n= 70, 33.8%), Lepidoptera (moths and butterflies; n= 65, 31.4%), and Hymenoptera (ants, bees, and wasps; n= 43, 20.8%). Coleoptera (beetles; n= 11, 5.3%) and Hemiptera (true bugs; n= 7, 3.4%) represent the next most frequently infected host orders. The remaining seven arthropod orders are each represented by only one or two species infected with Wmk-positive *Wolbachia* strains (Fig. 1D). Normalization of the count of Wmk-positive strains against the total number of *Wolbachia* strains within each host order revealed variation in prevalence across taxa. The highest prevalence was observed among strains infecting Coleoptera (100%), followed by Lepidoptera (~98%). It further declined in Diptera (86.4%), Hymenoptera (86%), and Hemiptera (~79%). Strains infecting coleopterans also exhibited the highest per-genome copy number of 2HTH-Wmk (median= 10; range= 1–18), comparable to that observed in orthopteran-associated strains (median= 10; range= 9–11) (Fig. 1F). However, these host orders are underrepresented in the dataset (Coleoptera: n= 11; Orthoptera: n= 2), which may inflate estimates of copy number variation due to limited sampling.

In contrast, among the three most represented host orders in the dataset, a median of four Wmk copies per *Wolbachia* genome was observed. The highest median copy number occurred in Lepidoptera-associated strains (median= 6; range= 1–22), followed by Hymenoptera (median= 4; range= 1–12) and Diptera (median= 3.5; range= 1–19) (Fig. 1F). Copy number variation in Lepidoptera-associated strains was statistically distinct from that in Diptera and Hymenoptera (Dunn’s test: unadjusted P< 0.0002; Bonferroni-adjusted P< 0.0008), indicating enrichment of Wmk copies in *Wolbachia* infecting moth and butterfly hosts. This pattern is consistent with previous observations based on comparative analysis of 32 *Wolbachia* genomes (Lefoulon et al., 2025).

### Wmk Type VI lineage

The comprehensive repertoire identified above revealed that 207 *Wolbachia* genomes collectively harbor 1205 non-redundant 2HTH-Wmk homologs. This dataset likely encompasses all previously recognized Wmk Types I–V (Lefoulon et al., 2025), as well as highly divergent variants resembling *w*Bif-Wmk and *w*Zbi-Wmk (Sahoo et al., 2026). Earlier analyses demonstrated that these divergent variants possess distinct sequence composition and reorganized structural features relative to canonical Type I Wmks (Sahoo et al., 2026). Building on these observations, the expanded dataset was used to test whether the divergent variants represent a discrete phylogenetic lineage or instead reflect independent, convergent sequence divergence within established Wmk types.

To address this question, phylogenetic trees were reconstructed using amino acid sequences from all 1205 homologs under maximum-likelihood (ML) and Bayesian inference (BI) frameworks. Using marker sequences delineated in Lefoulon et al. (2025), the majority of homologs (n= 1130) were confidently assigned to Wmk Types I–V (Fig. 2A; Fig. S1). These clades were strongly supported (BS and aLRT values > 99%; posterior probabilities [pp] > 0.9), consistent with the earlier phylogenetic reconstructions (Lefoulon et al., 2025). In contrast, 52 homologs formed a distinct, well-supported monophyletic clade (BS= 100%; aLRT= 100%; pp= 1) characterized by an extended basal branch (ML branch length= 0.8135). This basal branch was approximately 1.5–5.2 times longer than those subtending the five established Wmk type clades, indicating relatively higher evolutionary rate. This divergent lineage was therefore designated Wmk Type VI (Fig. 2A).

**Figure 2.**
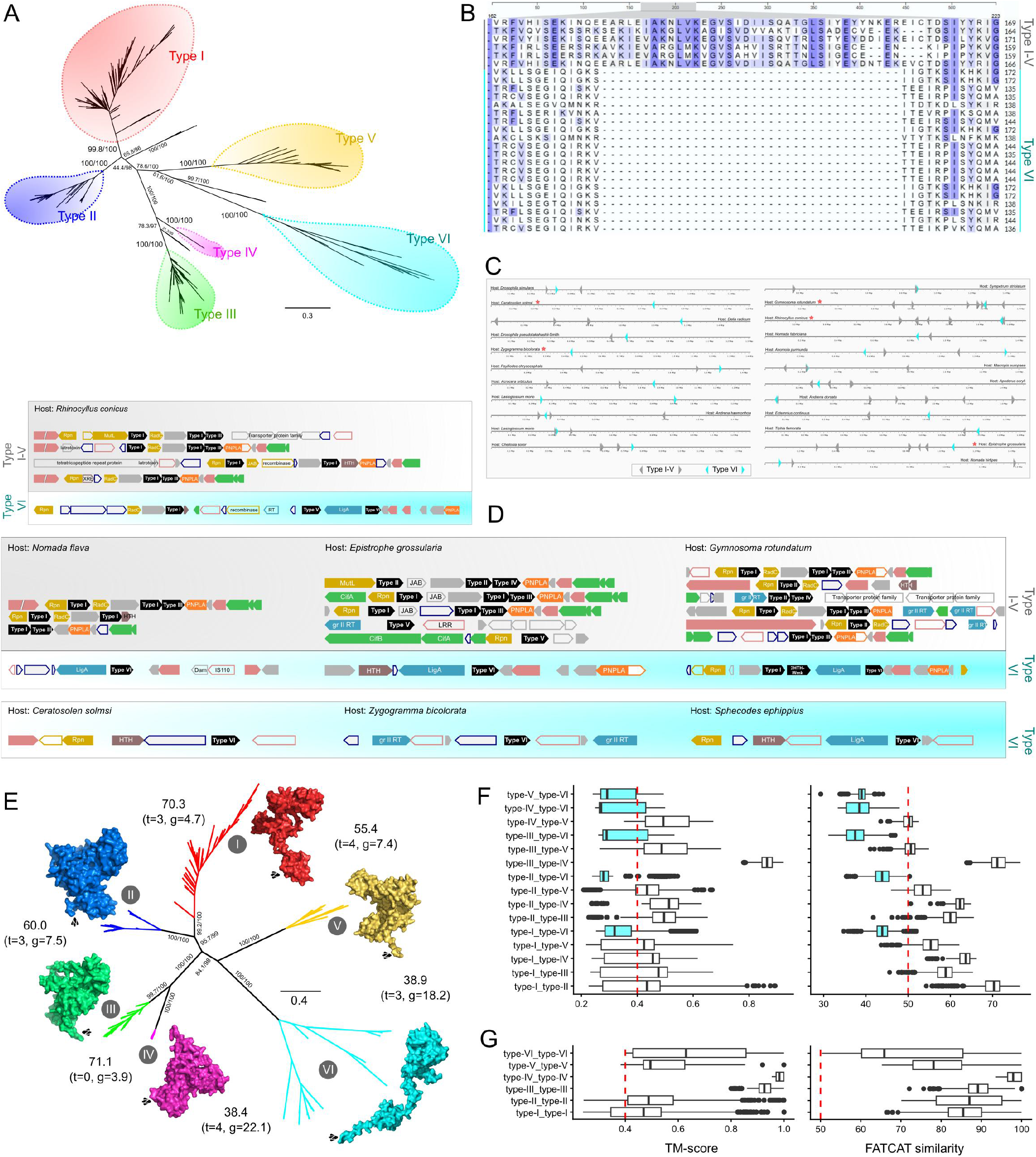
Phylogenetic and structural classification of Wmk homologs in *Wolbachia*. (A) Maximum likelihood (ML) phylogeny of 1205 Wmk homologs identified from 207 *Wolbachia* genomes. Node support values are shown as approximate likelihood ratio test and ultrafast bootstrap percentages (aLRT/BS) calculated from 1000 replicates. Major clades corresponding to Type I–VI groups are highlighted with colored balloons. Homologs not assigned to any of the six types are positioned outside the highlighted regions. (B) Representative alignment snapshot illustrating a characteristic sequence deletion in Type VI homologs (indicated by cyan side bars) relative to Type I–V homologs (indicated by gray side bars). Complete alignment file is provided in Fig. S2. (C) Genomic localization map of Wmk homologs in 23 selected *Wolbachia* strains with complete genome assemblies, chosen from the 34 genomes analyzed for neighborhood mapping. Genomes marked with an asterisk are also represented in panel (D). (D) Genomic neighborhood architecture of Wmk homologs in seven representative *Wolbachia* strains. Type VI homologs are shown with cyan background shading, whereas Type I–V homologs are depicted in gray. Flanking genes are included to illustrate local synteny patterns. Neighborhood maps for all 34 genomes are provided in Fig. S3. (E) Structure-based ML phylogeny of 166 Wmk homologs derived from 106 *Wolbachia* genomes. The tree was reconstructed using protein sequence and structural alphabets are separate partitions. Q3DiAF was used to model the structural partition, with node supports (aLRT/BS) based on 1000 replicates. Branches are colored according to sequence-based Type classification in panel (A) and labeled accordingly. Representative AlphaFold-predicted structures are displayed adjacent to their respective clades. Values at the tree periphery indicate median FATCAT similarity scores for all pairwise structural comparisons between the indicated left and right clades, with twists count and gap percentage shown in parentheses. A full table of FATCAT analysis is in Table S2. (F) Box plots summarizing the distribution of TM-scores and FATCAT similarity scores for all between-Type structural comparisons. (G) Box plots summarizing the distribution of TM-scores and FATCAT similarity scores for all within-Type structural comparisons.

An additional 23 homologs could not be assigned to any of the six defined types (I–VI). Instead, these sequences formed three minor but well-supported clades occupying distinct positions in the unrooted phylogeny (Fig. 2A; Fig. S1). A clade comprising 16 homologs (BS= 100%; aLRT= 100%; pp= 1) branched between Types I and II. A second clade of seven homologs (BS= 99%; aLRT= 31.3%; pp= 0.81) was positioned between Types III and IV. A third clade containing two homologs (BS= 100%; aLRT= 99.7%; pp= 1) diverged between Types V and VI. These lineages may represent currently unrecognized Wmk variant types or transitional forms arising from recurrent evolutionary trajectories. However, additional genomic sampling and functional characterization will be necessary to determine their phylogenetic and biological status.

The newly defined Type VI clade comprises 2HTH-Wmk homologs from *w*Bif, *w*Zbi, and *w*Pse N101, which were previously categorized as distant Wmk variants based on pronounced sequence divergence from the reference *w*Mel-Wmk and canonical Type I homologs (Sahoo et al., 2026). Comparative sequence analysis of Type VI homologs relative to Types I–V corroborated earlier observations and revealed the distinct molecular signature of the former: a deletion of ~36 amino acid residues upstream of the C-terminal HTH domain and an insertion of ~7 residues downstream of this domain (Fig. 2B; Fig. S2) (Sahoo et al., 2026). The conserved presence of these insertion-deletion features across all Type VI homologs is consistent with shared ancestry and provides a parsimonious explanation for their phylogenetic clustering. Importantly, removal of the alignment regions encompassing these characteristic insertion–deletion segments did not alter tree topology (Fig. S1). The stability of the clade following pruning indicates that Type VI monophyly is not an artifact of localized alignment differences but instead reflects gene-wide phylogenetic signal, thereby reinforcing inference of a common evolutionary origin.

### Genomic neighborhood analysis

Gene neighborhood analysis provided additional support for a distinct evolutionary trajectory of Type VI Wmk variants. To evaluate genomic context, locus maps encompassing up to six flanking genes on either side of each Wmk locus were compared across a representative set of 34 *Wolbachia* genomes spanning all six Wmk types (Fig. 2C, D; Fig. S3). Previously characterized neighborhood architectures from additional strains reported by Lefoulon et al. (2025) were incorporated for comparative assessment. Across Types I–V, Wmk homologs consistently localized within multi-gene cassettes enriched for DNA repair and recombination-associated functions (Lefoulon et al., 2025). These regions commonly encoded RadC, MutL, and members of the Rpn protein family, and were frequently associated with group II intron-encoded reverse transcriptases and ankyrin-repeat-containing proteins, although gene composition varied among strains. Consistent with prior observations (Perlmutter et al., 2019; Lefoulon et al., 2025), these Wmk-containing cassettes were often situated proximal to the WO prophage core region, as evidenced by neighboring genes encoding patatin-like phospholipases and phage structural components (Fig. 2D; Fig. S3).

In contrast, Type VI Wmk homologs were embedded within a markedly distinct genomic context. These loci were consistently associated with genes encoding DNA ligase and were frequently flanked by insertion sequence (IS) elements, with members of the IS3, IS5, IS110, and IS481 families frequently observed (Fig. 2D; Fig. S3). Additional neighboring genes included those encoding DNA adenine methylase and reverse transcriptases. Collectively, this neighborhood architecture differs substantially from the canonical Type I–V Wmk cassettes. Despite these structural differences, the gene complements of both neighborhood types suggest potential functional convergence. The presence of repeat elements in Type I–V loci and IS elements in Type VI neighborhoods indicates elevated genomic mobility in both contexts. Moreover, enrichment of DNA repair-associated genes in Type I–V cassettes and the presence of DNA ligase in Type VI neighborhoods point toward a shared role in maintaining DNA integrity.

Notably, Type VI homologs were frequently observed as isolated loci within their immediate genomic context (Fig. 2C, D; Fig. S3). However, in several *Wolbachia* genomes, Type VI variants occurred in close proximity to Type I or Type V homologs. For example, in strains infecting *Rhinocyllus conicus, Macropis europaea*, and *Andrena dorsata*, Type VI and Type V homologs were separated by a single gene encoding DNA ligase (Fig. 2C; Fig. S3). Similarly, in the strain infecting *Gymnosoma rotundatum*, the Type VI homolog was positioned adjacent to a Type I Wmk variant (Fig. 2C, D). Importantly, the characteristic association between Type VI homologs and DNA ligase was preserved across these structural rearrangements, although the ligase gene was pseudogenized in multiple instances.

### Structure based comparisons

The unique sequence composition of Type VI homologs is likely associated with structural remodeling (Sahoo et al., 2026). Representative sequence comparisons have shown that, relative to Type I variants, Type VI homologs possess a markedly shorter inter-domain linker, a feature predicted to induce global conformational differences between the two types (Sahoo et al., 2026). Given that protein tertiary structures are generally more conserved over evolutionary timescales than primary amino acid sequences (Illergard et al., 2009), evolutionary divergence of Type VI homologs was further examined using structure-based phylogenetic analyses. For this purpose, AlphaFold-predicted protein structures of 166 selected homologs derived from 106 *Wolbachia* genomes were analyzed (Fig. S4). This dataset included representatives of all six Wmk types, comprising the complete set of 52 Type VI variants and a randomly sampled subset of 114 homologs representing Types I–V.

For maximum-likelihood–based phylogenetic reconstruction of protein 3D folds, a recently benchmarked partitioned framework was applied in which sequence- and structure-derived features are jointly incorporated into a single inference while modeled as independent partitions (Mutti et al., 2025). In this framework, evolutionary variation in the sequence partition was modeled using the GTR substitution matrix, whereas structural variation was modeled using either the 3DiPhy (Puente-Lelievre et al., 2023) or Q3DiAF (Garg and Hochberg, 2025) substitution matrix in two separate analyses. Phylogenetic trees inferred under both structure-aware models consistently recovered all six Wmk types with strong statistical support (BS > 99; aLRT > 99) (Fig. 2E; Fig. S5). Further evaluation of phylogenetic relationships was conducted using model-independent approach implemented in FoldTree (Moi et al., 2025). In this approach, tree is reconstructed from statistically corrected distance matrix derived from structure-guided alignments integrating 3Di structural alphabets with amino acid sequences (van Kempen et al., 2024) (Fig. S5). These structure-informed phylogenies were topologically congruent with the sequence-only reconstructions, as suggested by the deep node divergence patterns, reflecting minimal discordance across analytical frameworks, and robustly supporting the delineation of Wmk proteins into six distinct phylogenetic types.

To assess global structural differences among the phylogenetic groups, predicted structures were compared using two independent approaches. First, TM-scores were calculated from pairwise structural alignments generated using the combined 3Di+AA algorithm implemented in FoldSeek (van Kempen et al., 2024). Second, pairwise structural similarity was estimated using flexibility-aware alignments implemented in FATCAT (Li et al., 2020). Whereas the former approach integrates structural information within a sequence-alignment framework, the latter explicitly accommodates conformational variability through twists and flexible gaps, rendering it well suited for modular and structurally dynamic proteins such as Wmk (Sahoo et al., 2026). Across Types I–V, global structural configurations remained broadly conserved despite phylogenetic divergence, as indicated by between-type comparisons showing median TM-scores >0.4 and median FATCAT similarity values >50% (Fig. 2F; Table S2). In certain comparisons – particularly between Types I and II – the TM-score distribution included values below 0.4, consistent with increased structural dispersion (Fig. 2F). However, this pattern largely reflects substantial within-type structural variability (Fig. 2G), likely arising from the modular architecture and conformational flexibility characteristic of Wmk homologs (Sahoo et al., 2026), as well as the reliance on static structural models in the present analysis.

In contrast, Type VI homologs exhibited high within-type structural similarity (median TM-score >0.4; median FATCAT similarity >50%) (Fig. 2G), indicating conservation of global fold among members of this clade. However, comparisons between Type VI and Types I–V revealed pronounced structural divergence, with median TM-scores ranging from 0.27 to 0.32 and median FATCAT similarity values between 37% and 44% (Fig. 2F; Table S2). These values fall substantially below those observed in inter-type comparisons among Types I–V, supporting a marked shift in global structural configuration associated with the Type VI lineage. Notably, pairwise comparisons between Type VI and Type I-V – particularly Types I and II – also included TM-scores exceeding 0.4 (Fig. 2F), indicating partial structural similarity in a subset of comparisons. This dispersion likely attributable to the intrinsic conformational flexibility of Wmk as noted above, or to the presence of underlying structural polymorphism shared among Types I, II, and VI, which needs further investigation.

### Wmk distribution across Wolbachia

To investigate macro-evolutionary correlates associated with the distinctive molecular features of the Type VI Wmk variant, its distribution was examined across *Wolbachia* supergroups and host taxonomic orders and compared with the patterns observed for Types I–V. For this analysis, 1182 Wmk homologs – excluding sequences not assignable to Types I–VI – were mapped across 203 *Wolbachia* genomes and projected onto a global *Wolbachia* phylogeny reconstructed from 251 whole genomes (Fig. 3A; Fig. S6).

**Figure 3.**
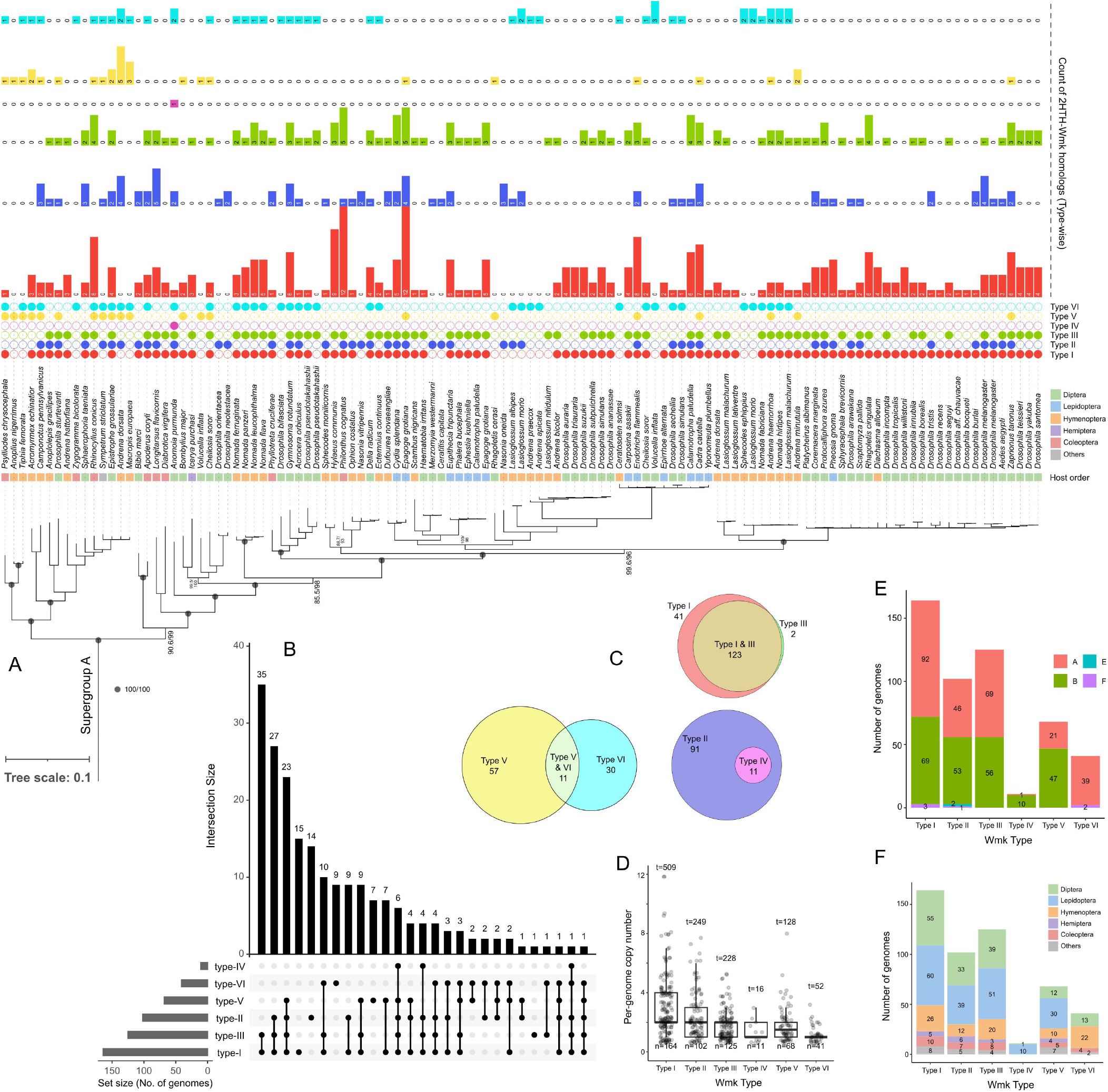
Phylogenetic distribution and copy number variation of Wmk homologs across *Wolbachia* genomes. (A) Maximum likelihood (ML) phylogeny of Supergroup A *Wolbachia*, pruned from the global phylogeny reconstructed using 251 genomes. Other clades of the global phylogeny is provided in Fig. S6. Node support values are shown as approximate likelihood ratio test and ultrafast bootstrap percentages (aLRT/BS) based on 1000 replicates; support values are displayed for deeper nodes only. Tip labels indicate host species name, host taxonomic order, and the presence and copy number of full-length Wmk homologs spanning Type I–VI. (B) UpSet plot depicting genome-wise distribution of Wmk homologs across Type I–VI for 203 *Wolbachia* genomes harboring a total of 1182 homologs. (C) Venn diagrams illustrating the co-occurrence of specific Wmk Type combinations across genomes, with numbers indicating the count of genomes harboring each combination or Type. (D) Box plot illustrating per-genome copy number variation across all Wmk Types. For each Type, the total number of genomes harboring at least one homolog (n) is shown at the bottom of the plot, while the cumulative number of homolog copies across those genomes (t) is indicated at the top. (E) Distribution of Type I–VI Wmk homologs across 203 *Wolbachia* genomes, grouped by supergroup classification. (F) Distribution of Type I–VI Wmk homologs across the same 203 genomes, categorized by host taxonomic order.

The distribution of Wmk types across *Wolbachia* genomes was markedly uneven. Type I was the most prevalent, comprising ~43% of all homologs (509/1182) and occurring in ~81% of genomes (164/203) (Fig. 3A, B). In contrast, the remaining types exhibited more restricted distributions, ranging from Type IV, detected in only ~5% of genomes (11/203), to Type III, present in ~62% (125/203). Type VI showed moderate prevalence, occurring in ~20% of genomes (41/203) (Fig. 3A, B). This heterogeneity reflects variation in both genome-level occurrence and within-genome copy-number: multiple Wmk types frequently co-occurred within individual genomes, and several genomes contained multiple copies of the same type. Notably, eight *Wolbachia* genomes encoded homologs encompassing five Wmk types (Fig. 3B). In two genomes, the Type I variant underwent pronounced expansion, reaching 12 copies each – the highest type-specific amplification observed (Fig. 3A, D). In comparison, Types II–V attained a maximum of 3–8 copies per genome, whereas Type VI homologs were limited to at most three copies per genome with the median value of one (Fig. 3D).

A notable feature of Type VI was its phylogenetic restriction. Unlike Types I–V, which were distributed across both A- and B-supergroup *Wolbachia*, Type VI homologs were detected almost exclusively within the A-supergroup, with only two occurrences in Supergroup F (Fig. 3E; Fig. S6). Within the A-supergroup, Type VI was present in ~33% of strains (39/117), whereas Types I–V collectively occurred in ~93% (109/117). In nine strains – including the well-characterized *w*Zbi (host: *Zygogramma bicolorata*) and *w*Bif (host: *Drosophila bifasciata*) – Type VI appeared in isolation, without detectable co-occurrence of other Wmk types (Fig. 3A; Fig. S3). However, the presence of additional truncated or single-HTH variants cannot be excluded; for instance, a Type III 1HTH-Wmk was previously reported in *w*Bif (Lefoulon et al., 2025).

In addition to phylogenetic confinement, Type VI exhibited a pronounced host-order bias. Approximately 85% of Type VI occurrences (35/41) were associated with *Wolbachia* infecting Hymenoptera and Diptera (Fig. 3F). By contrast, these two host orders accounted for only ~53% (102/194) of strains harboring Types I–V, whereas Lepidoptera contributed ~33% (65/194). Notably, Type VI homologs were entirely absent from *Wolbachia* infecting Lepidoptera (Fig. 3F), despite strong representation of this host order in the dataset (~32%; 65/203). This absence is particularly striking given that Lepidoptera-associated *Wolbachia* are known to encode additional male-killing determinants distinct from Wmk (Katsuma et al., 2022). Collectively, these findings indicate that Type VI follows a comparatively constrained evolutionary trajectory, characterized by restricted supergroup distribution, limited copy-number expansion, and marked host-order specificity, distinguishing it from other Wmk types.

## Conclusion

Patterns of effector gene diversity are shaped by multiple, interacting mechanisms, including sequence divergence, presence/absence polymorphism, copy number variation, and regulatory differences (Muller et al., 2026). Recent advances in structural genomics, particularly those driven by structure-predicting large language models, have further demonstrated that structural variation contributes substantially to effector diversification beyond primary sequence changes alone (Moi et al., 2025; Seong and Krasileva, 2023). The analyses presented here indicate that diversity of the full-length Wmk effector in *Wolbachia* reflects the combined influence of these processes. First, sequence diversity pattern largely corroborated structural diversity revealing six phylogenetic types of Wmk variants. This finding reinforce the five previously recognized lineages (Lefoulon et al., 2025) and additionally identify a sixth, highly divergent lineage (Type VI). Type VI homologs are characterized by extensive sequence divergence and marked structural reorganization relative to Types I–V, consistent with a distinct evolutionary trajectory. The functional implications of this reorganization – whether involving loss, gain, or modification of activity – remain to be experimentally evaluated. However, the genomic context of Type VI differs consistently from that of other lineages, supporting its unique evolutionary history and suggesting potential differences in regulatory control.

Next, strain-level polymorphism in presence/absence patterns and copy number variation is pronounced across Wmk lineages. Type I variants occur in approximately 81% of surveyed *Wolbachia* genomes and exhibit the highest per-genome copy number (Fig. 3D). Such widespread distribution and dosage enrichment are consistent with the interpretation that Type I represents a core component of the Wmk-associated phenotype, in agreement with previous evidence linking male killing to the presence or expression of this variant (Arai et al., 2023; Permutter et al., 2021, 2019). In contrast, Type III has been shown to modulate the phenotypic effect of Type I (Lefoulon et al., 2025; Arai et al., 2023), indicating that diversification within the Wmk repertoire contributes to quantitative or qualitative fine-tuning of the male-killing phenotype rather than acting solely through binary gain/loss dynamics. Consistent with this interpretation, Type I and Type III co-occur in 123 genomes, frequently in tandem configuration (18/23 cases with resolved local arrangement) (Fig. 3C; Fig. S3), corroborating previous observation (Lefoulon et al., 2025). The recurrent physical association of these variants suggests selective maintenance of linkage and raises the possibility that they function as a coordinated module, representing a coupled functional and evolutionary unit within the Wmk effector repertoire.

In contrast to the broad distribution of Type I, the Type VI variant is comparatively rare, occurring in 16% of surveyed *Wolbachia* genomes and predominantly as one or two copies per genome (Fig. 3D). This lineage exhibits a restricted phylogenetic distribution, being largely confined to Supergroup A *Wolbachia*, with only two reported occurrences in Supergroup F, and is notably absent from strains associated with Lepidoptera hosts (Fig. 3E, F). The observed distributional bias is unlikely to be attributable to uneven genome sampling, as the 251-genome dataset included substantial representation across supergroups, with approximately 31% of strains derived from Lepidoptera-associated *Wolbachia* (Fig. 1B). In the absence of empirical data linking Type VI to specific phenotypic outcomes, the evolutionary basis of its restricted occurrence remains unresolved. Nevertheless, the present/absence polymorphism of the Type VI Wmk variant appears to be structured by *Wolbachia* phylogenetic delineation and host association patterns. A comparatively recent evolutionary origin may also contribute to its limited taxonomic spread, although this hypothesis requires further temporal resolution.

Beyond the six Wmk types resolved in the present analysis, additional variants remain either unclassified within the current framework (Fig. 2A) or likely remain undiscovered, indicating that the diversity of this effector exceeds what is presently captured. The scale and structural organization of this variation are inconsistent with a simple model of sequential allelic replacement driven exclusively by directional host-symbiont arms race dynamics. Although reciprocal evolutionary interactions between *Wolbachia* and its hosts – particularly under gene-for-gene–like scenarios – may contribute to diversification, the recurrent co-occurrence of multiple Wmk variants within single genomes argues against a model dominated by recurrent selective sweeps of single advantageous alleles. Instead, the observed patterns are more consistent with functional diversification mediated by context-dependent modulation. Wmk variants may retain overlapping mechanistic roles while differing in regulatory architecture, expression timing, interaction specificity, or functional efficiency within distinct host cellular environments. Such divergence could involve differential perturbation of host sex-determination pathways – for example, through effects on dosage compensation – or adaptation to lineage-specific translational and biochemical contexts, including codon usage compatibility (Permutter et al., 2021). Within this framework, diversification does not necessarily entail acquisition of entirely novel molecular functions, but rather refinement and optimization of effector performance across heterogeneous host backgrounds.

Given the broad host range of *Wolbachia* and its documented capacity for horizontal transmission (Vancaester and Blaxter, 2023; Scholz et al., 2020), maintenance of multiple effector variants may provide standing genetic variation that facilitates establishment and persistence in novel host lineages. This scenario is more consistent with trench-warfare–like dynamics, characterized by long-term maintenance of allelic diversity, than with repeated hard selective sweeps (Sanchez-Vallet et al., 2018). Although functional validation will be required to distinguish among these evolutionary models, the distribution, coexistence, and structural divergence of Wmk types collectively support an interpretation in which diversification reflects adaptive flexibility rather than simple antagonistic escalation.

## Methods

### Identifying Wmk homologs

A comprehensive dataset of *Wolbachia* genomes was assembled by combining genomes analyzed previously in Sahoo et al. (2025) and Lefoulon et al. (2025), resulting a total of 251 genomes. Genome redundancy was minimized by retaining a single representative for each unique combination of *Wolbachia* supergroup, host species, and phylogenetic relatedness. Specifically, when multiple closely related strains belonged to the same supergroup and were isolated from the same host species, only one strain was retained. All genomes were annotated de novo for protein-coding sequences using Prokka v1.14.6 (Seemann 2014). Wmk loci were identified using three previously characterized Wmk proteins as references: *w*Mel-Wmk (AAS14326), *w*Bif-Wmk (QEQ51101), and *w*Zbi-Wmk (XXB89297). Homology detection was performed using MMseqs2 v16.747c6 (Steinegger and Söding 2017), with a minimum sequence identity threshold of 40% to mitigate false negatives arising from sequence divergence near the “twilight zone.”

Putative homologs were subsequently filtered based on alignment coverage relative to the reference sequences. Only sequences whose alignments spanned both the N-terminal and C-terminal helix–turn– helix (HTH) regions of the references were retained, thereby restricting the dataset to full-length Wmk variants containing two HTH domains (2-HTH). When a single query sequence aligned to multiple reference Wmk variants, the hit with the highest alignment score was selected. To validate that the retained sequences indeed encode HTH domains, a representative subset of 102 homologs was analyzed against the Prosite database (Sigrist et al., 2013) using the ScanProsite online portal (de Castro et al., 2006).

### Phylogenetic reconstruction

The phylogeny of 251 *Wolbachia* genomes was reconstructed using shared BUSCO gene markers, following the approach described in Sahoo et al. (2025). All genomes were screened for conserved single-copy orthologs against the Rickettsiales_odb10 database (number of BUSCOs: 345) using BUSCO v6.0 (Manni et al., 2021). A set of 323 BUSCO markers present in at least 95% of the genomes was selected for downstream analyses. These markers were concatenated into a supermatrix alignment using the BUSCO phylogenomics pipeline v20240919 (github.com/jamiemcg/BUSCO_phylogenomics). The use of BUSCO markers ensured that phylogenetic inference relied on core genomic elements that are largely devoid of prophage and repetitive regions, which often exhibit evolutionary histories distinct from the bacterial core genome. The resulting alignment matrix was used to infer a maximum-likelihood (ML) phylogeny in IQ-TREE v2.1.4 (Minh et al., 2020) under the LG+G4 substitution model. Node support was assessed using 1000 ultrafast bootstrap replicates and the approximate likelihood ratio test (aLRT).

For Wmk-specific phylogenetic analyses, all 1205 homologs identified across the 251 genomes were aligned using MAFFT v7.490 (-auto) (Katoh and Standley 2013) and subsequently trimmed to correct for alignment errors using trimAl v1.5 (-automated1) (Capella-Gutiérrez et al., 2009). The curated alignment was used for ML tree reconstruction in IQ-TREE v2.1.4 (Minh et al., 2020), with the best-fit evolutionary model selected using ModelFinder. Branch support values were estimated from 1000 bootstrap replicates and aLRT statistics. Bayesian phylogenetic inference was conducted on the same trimAl-corrected alignment using MrBayes v3.2.7 (Ronquist et al., 2012), with two independent runs of five million generations, sampling every 500 generations, and discarding the first 25% of samples as burn-in. In an independent analysis, the 1205 Wmk homologs identified in this study were combined with 109 previously characterized homologs from Lefoulon et al. (2025) and analyzed using the same ML framework described above. Previously defined Wmk types from Lefoulon et al. (2025) were used to assign type designations to homologs in the current dataset.

### Gene neighborhood analysis

To characterize the genomic neighborhoods of Wmk homologs and generate comparative visualizations, gene context analyses were performed using the online tool webFlaGs (Saha et al., 2021). As webFlaGs relies on RefSeq-based annotations for neighborhood mapping, a subset of genomes was reanalyzed using RefSeq annotations rather than the Prokka annotations applied to the primary dataset.

A curated set of 34 *Wolbachia* genomes harboring Type VI Wmk homologs was selected for this analysis. These genomes were verified to also contain additional Wmk homologs belonging to other types (Types I–V). Genome assemblies, annotations, and protein sequences were retrieved from the RefSeq database (O’Leary et al., 2016) and screened for Wmk homologs using the same homology detection criteria described above. The corresponding genome accessions and Wmk protein identifiers were uploaded to webFlaGs to extract gene neighborhoods spanning up to six flanking genes on either side of the Wmk locus. The resulting neighborhood maps were manually inspected against RefSeq annotation files to verify gene identities and were subsequently color-coded for visualization.

To reassign Wmk types to homologs derived from the 34 RefSeq genomes, these sequences were phylogenetically placed onto the previously inferred Wmk tree comprising 1205 homologs. Specifically, RefSeq-derived homologs were combined with the original dataset, aligned using MAFFT v7.490 (Katoh and Standley 2013), and trimmed with trimAl v1.5 (Capella-Gutiérrez et al., 2009). Maximum-likelihood phylogenetic inference was performed using the original Wmk tree as a constraint in IQ-TREE v2.1.4 (Minh et al., 2020). Wmk type assignments for the newly added homologs were determined based on their placement within previously defined type-specific clades.

### Protein structure analysis

A total of 166 Wmk homologs were randomly selected for protein structure analyses, collectively representing all six Wmk types. 3D structural models were predicted using ColabFold v1.5.5 (Mirdita et al., 2022), an implementation of AlphaFold2 (AF2) (Jumper et al., 2021). Predictions were generated via the Colab notebook using default parameters. Model quality and structural features were evaluated using confident matrices from AF2 and visualized with PyMOL v2.6 (open-source build). For downstream structure-based analyses, the top-ranked AF2 model for each homolog was used.

Structure-based phylogenetic analyses was conducted using three different tree reconstruction strategies that differed in alignment representation and evolutionary models. All alignments were generated using MAFFT v7.490 (Katoh and Standley 2013) and trimmed with trimAl v1.5 (Capella-Gutiérrez et al., 2009). As a reference, amino acid sequences corresponding to the 166 modeled proteins were aligned and analyzed under the LG (F+G4) substitution model. For FoldTree analysis (Moi et al., 2025), structures were analyzed using Colab notebook in default settings. For model based analysis, structures were first encoded as 3Di alphabets using FoldSeek v9.427 (van Kempen et al., 2024). Then, both amino acid and 3Di alignments of structures were combined and treated as separate partitions. In these analyses, the amino acid partition was modeled using LG (+F+G4), while the 3Di partition was modeled using either 3DiPhy (+F+G4) or Q3DiAF (+F+G4). All maximum-likelihood analyses were conducted in IQ-TREE v2.1.4 (Minh et al., 2020), with node support assessed using bootstrap resampling and the approximate likelihood ratio test.

## Acknowledgements

The work was supported by the DST-INSPIRE Faculty Fellowship to RKS (award no. DST/INSPIRE/04/2019/000478). RKS thanks the high-performance computing facility at CSIR-CCMB, Hyderabad. RKS thanks Karthikeyan Vasudevan for academic support during the execution of this work.

## Data availability

No additional data generated during the study.

