## Supplementary material for "Structural reorganization and genomic context define a divergent lineage of the *Wolbachia* male-killing gene *wmk*": Tables S1-S2 and Figures S1-S6

**Supplementary file** for the title: Structural reorganization and genomic context define a divergent lineage of the *Wolbachia* male-killing gene *wmk*

**Table S1.** Comparative summary of full-length Wmk homolog counts in 30 *Wolbachia* strains, contrasting the numbers detected in the present study with those reported in a previous investigation.

| Strain Name | Genome accession | GenBank accession | Count of 2HTH-Wmk in current study | Count of 2HTH-Wmk in Lefoulon et al 2025 |
|---|---|---|---|---|
| *w*Ana | GCA_008033215 | CP042904 | 3 | 3 |
| *w*Biau1 | GCA_037076435 | AP028655 | 0 | 0 |
| *w*Biau2 | GCA_037076445 | AP028656 | 0 | 1 |
| *w*Bif | GCA_014129685 | JAATLC010000010 | 1 | 2 |
| *w*Bol1 | GCA_000333775 | CAOH01000062 | 5 | 5 |
| *w*Bor | GCA_014129615 | JAATLD010000002 | 3 | 3 |
| *w*CauA | GCA_045865005 | AP028948 | 14 | 17 |
| *w*CauB | GCA_045865015 | AP028949 | 11 | 12 |
| *w*CI | GCA_045865035 | AP028951 | 4 | 6 |
| *w*Fur | GCA_023559125 | CP096925 | 6 | 7 |
| *w*Ha | GCA_000376605 | CP003884 | 3 | 3 |
| *w*Hm-t | GCA_030295095 | AP025638 | 7 | 8 |
| *w*Inn | GCA_021378375 | CP076228 | 3 | 3 |
| *w*Kue | GCA_045865025 | AP028950 | 2 | 2 |
| *w*Mau | GCA_004795975 | CP034335 | 1 | 2 |
| *w*Mel | GCA_000008025 | AE017196 | 4 | 5 |
| *w*No | GCA_000376585 | CP003883 | 1 | 1 |
| *w*Pip | GCA_000073005 | NC_010981 | 4 | 4 |
| *w*Pse-N101 | GCA_026274225 | JAPJVH010000002 | 3 | 4 |
| *w*Pse-Smith | GCA_026015925 | CP110359 | 3 | 4 |
| *w*Rec | GCA_000742435 | JQAM01000018 | 1 | 1 |
| *w*Ri | GCA_000022285 | CP001391 | 6 | 6 |
| *w*Sca | GCF_023559145 | CP096926 | 6 | 7 |
| *w*Spc | GCA_002300525 | NTHL01000073 | 3 | 3 |
| *w*Tpre | GCA_001439985 | CM003641 | 0 | 0 |
| *w*VulC | GCA_041439025 | CP156068 | 9 | 8 |
| *w*VulM | GCA_041440575 | CP156069 | 5 | 6 |
| *w*VulP | GCA_041442225 | CP156070 | 6 | 6 |
| *w*Wil | GCA_040084705 | CP157591 | 4 | 4 |
| *w*Yak | GCA_018467115 | CP069053 | 6 | 6 |

**Table S2.** Median structural similarity metrics among the six Wmk Types based on pairwise conformational comparisons. FATCAT scores are presented in the lower-left panel of the matrix, whereas TM-align TM-scores are shown in the upper-right panel. Cyan-colored cells denote comparisons involving Type VI against other Types.

| | Type I | Type II | Type III | Type IV | Type V | Type VI |
|---|---|---|---|---|---|---|
| Type I | | 0.434 (55.00) | 0.475 (42.00) | 0.455 (42.90) | 0.424 (37.10) | 0.318 (27.20) |
| Type II | 70.315 (3, 4.68) | | 0.495 (42.20) | 0.513 (44.55) | 0.434 (37.70) | 0.278 (26.90) |
| Type III | 58.88 (4, 7.17) | 60 (3, 7.54) | | 0.864 (56.10) | 0.487 (34.80) | 0.290 (24.70) |
| Type IV | 63.64 (3, 5.66) | 62.26 (3, 5.98) | 71.1 (0, 3.9) | | 0.493 (32.70) | 0.270 (24.90) |
| Type V | 55.35 (4, 7.42) | 53.47 (3, 6.93) | 50.65 (4, 9.52) | 50.24 (3, 8.14) | | 0.292 (23.50) |
| Type VI | 43.88 (3, 17.88) | 43.84 (3, 17.82) | 37.46 (4, 22.3) | 38.44 (4, 22.07) | 38.99 (3, 18.18) | |

TM-score (pident)

FATCAT similarity (twists count, gaps percent)

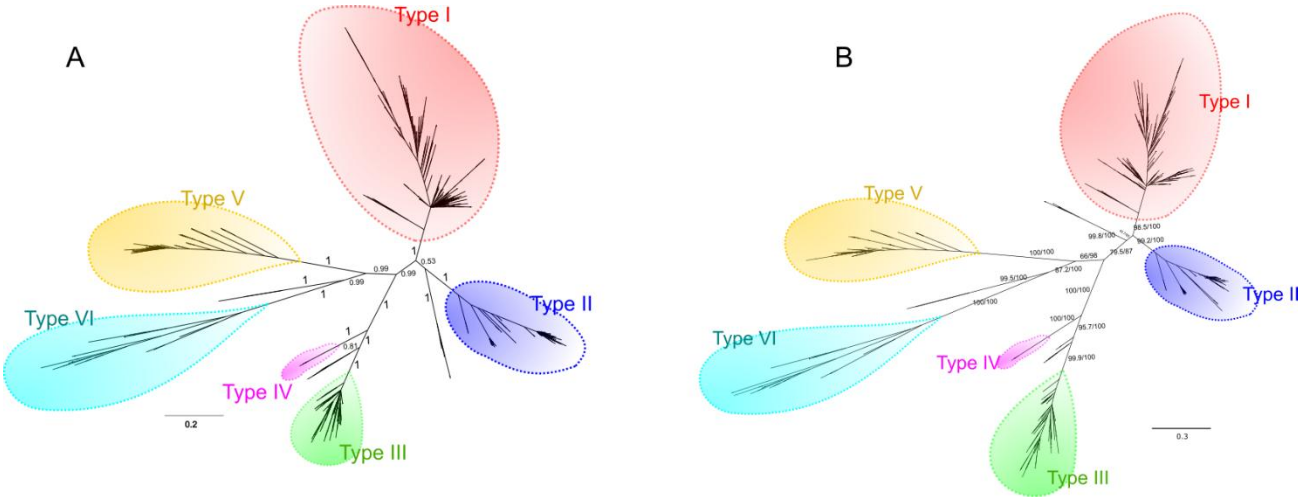


**Figure S1.** Phylogenetic reconstruction of Wmk homologs under alternative inference frameworks. (A) Bayesian phylogeny of 1205 Wmk homologs identified from 207 *Wolbachia* genomes. Node support values are posterior probabilities and are shown for deeper nodes only. Major clades corresponding to Type I–VI groups are highlighted with colored balloons. Homologs not assigned to any of the six types are positioned outside the highlighted regions.

(B) Maximum likelihood phylogeny of the same 1205 Wmk homologs after pruning the alignment to remove positions 99–145, which encompass the characteristic deletion observed in Type VI relative to Type I–V. Node support values are shown as approximate likelihood ratio test and ultrafast bootstrap percentages (aLRT/BS) calculated from 1000 replicates. Major clades corresponding to Type I–VI groups are highlighted with colored balloons, and unclassified homologs are positioned outside these regions.

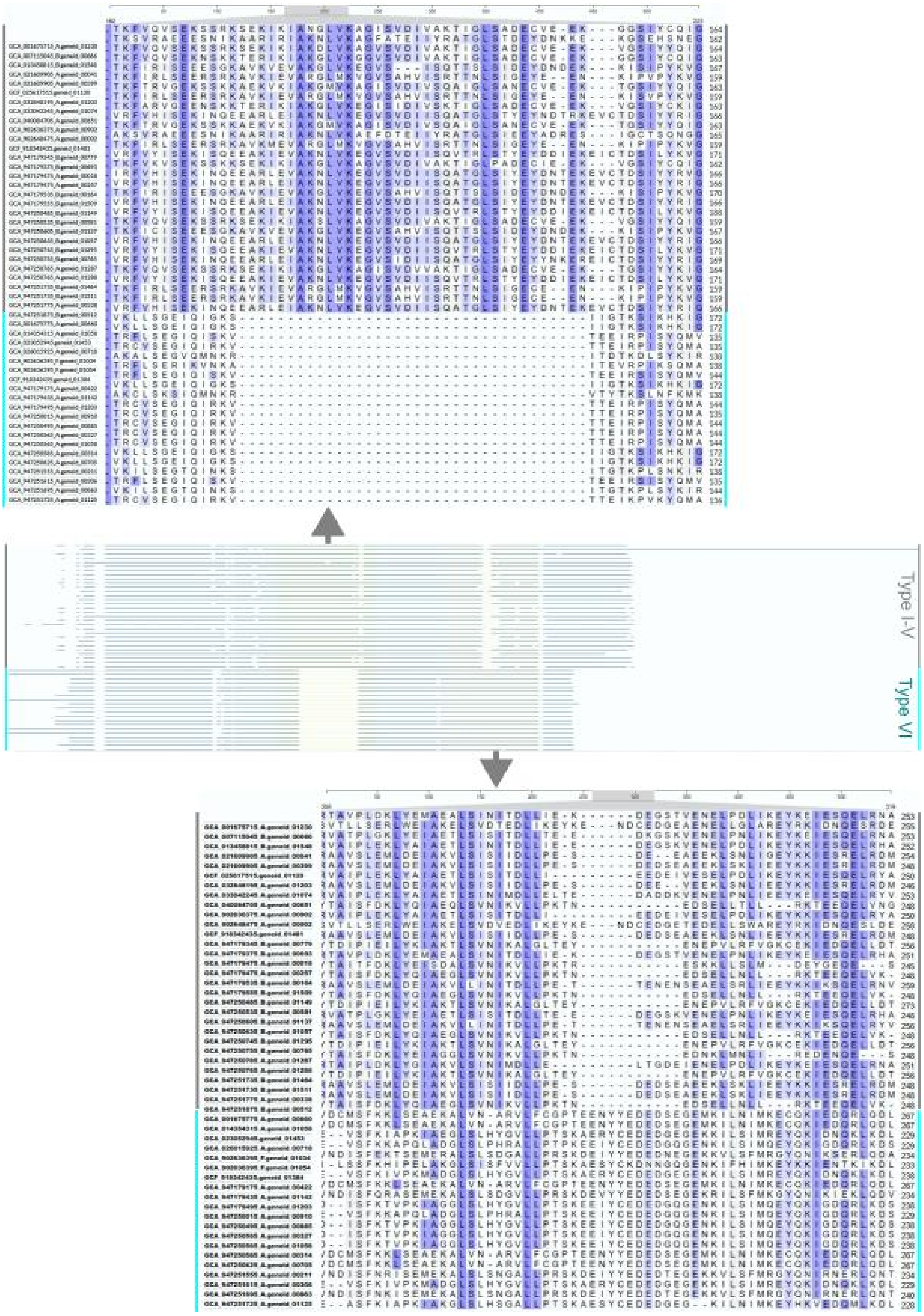


**Figure S2.** Snapshot of the multiple sequence alignment of 50 Wmk homologs highlighting the region encompassing the characteristic sequence deletion (upper panel) and insertion (lower panel) in Type VI homologs. The alignment was generated using Clustal Omega via the UniProt server and includes 30 Type I–V homologs (indicated by gray side bars) and 20 Type VI homologs (indicated by cyan side bars).

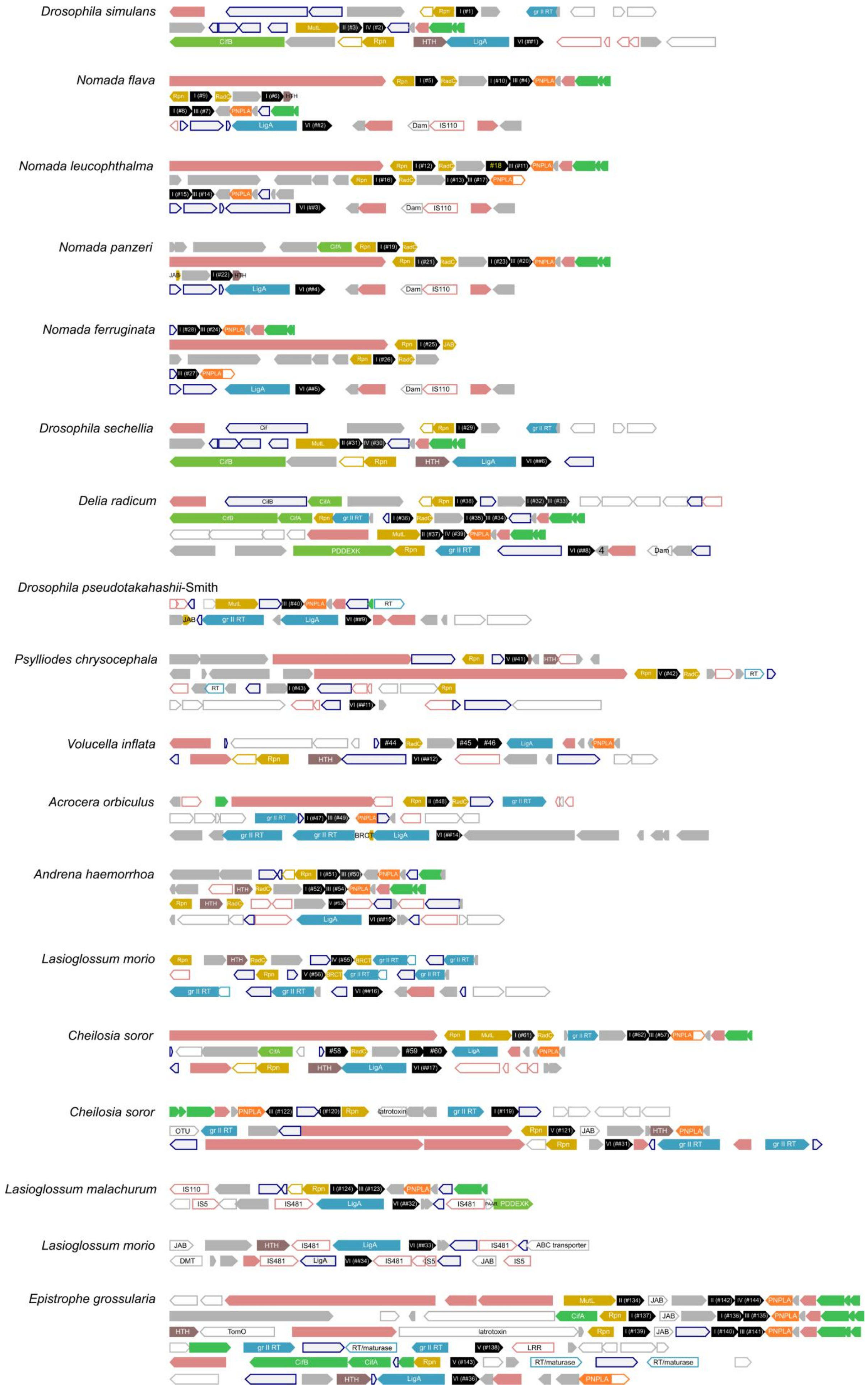
Drosophila simulans
Nomada flava
Nomada leucophthalma
Nomada panzeri
Nomada ferruginata
Drosophila sechellia
Delia radicum
Drosophila pseudotakahashii-Smith
Psylliodes chrysocephala
Volucella inflata
Acrocera orbiculus
Andrena haemorrhoa
Lasioglossum morio
Cheilosia soror
Cheilosia soror
Lasioglossum malachurum
Lasioglossum morio
Epistrophe grossularia

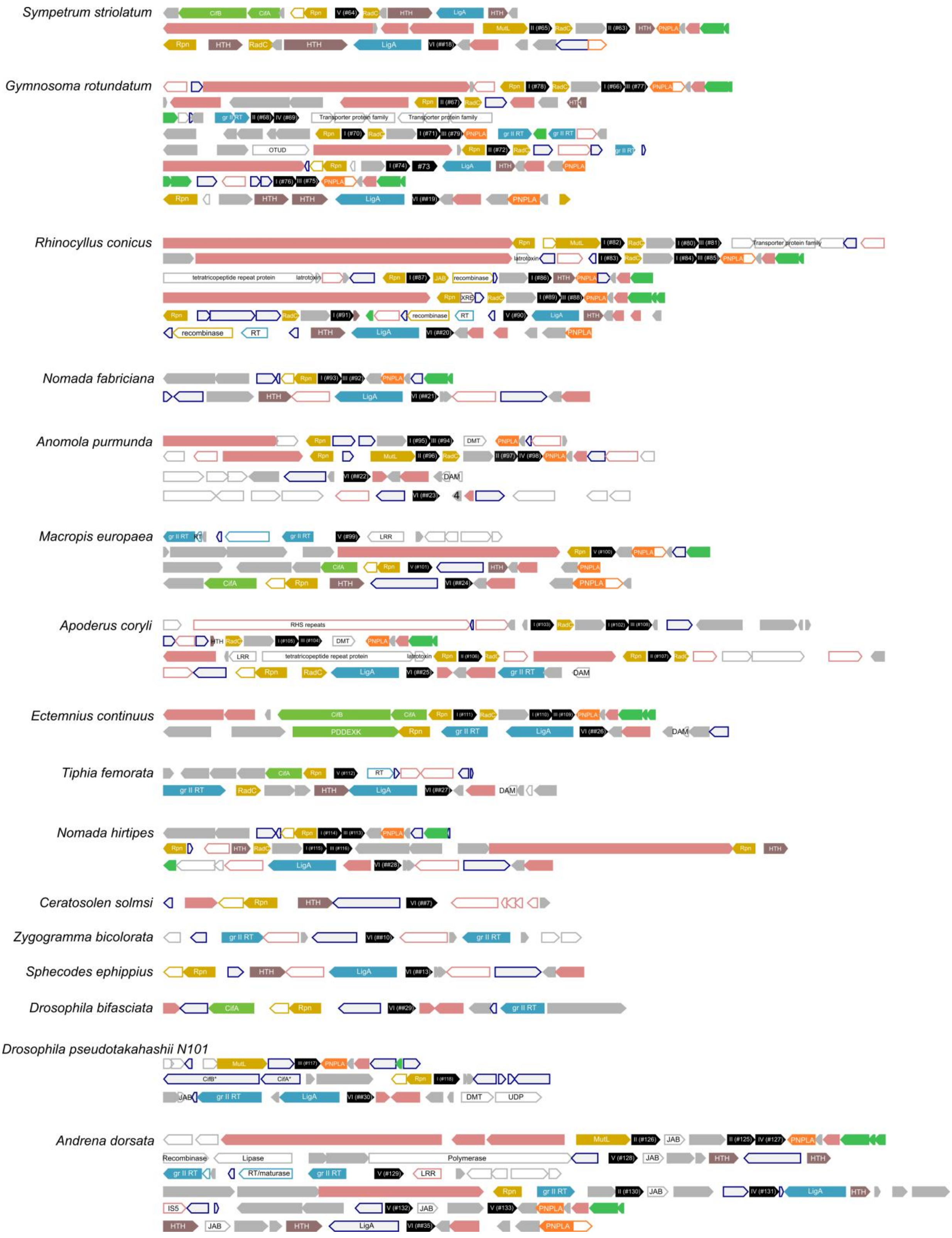


**Figure S3.** Genomic neighborhood maps of Wmk homologs in 34 *Wolbachia* genomes. Flanking genes are shown to illustrate local synteny patterns.

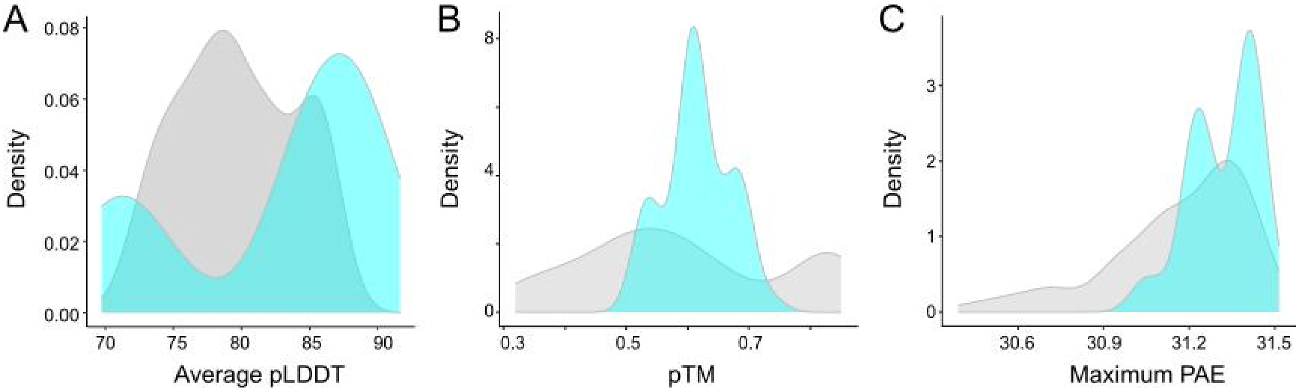


**Figure S4.** Distribution of model confidence metrics for AlphaFold-predicted structures of 166 Wmk homologs. Shown are the distributions of average pLDDT, predicted TM-score (pTM), and maximum predicted aligned error (max PAE). Values for Type VI homologs (52 structures) are indicated in cyan, whereas those for Type I–V homologs (114 structures) are shown in gray.

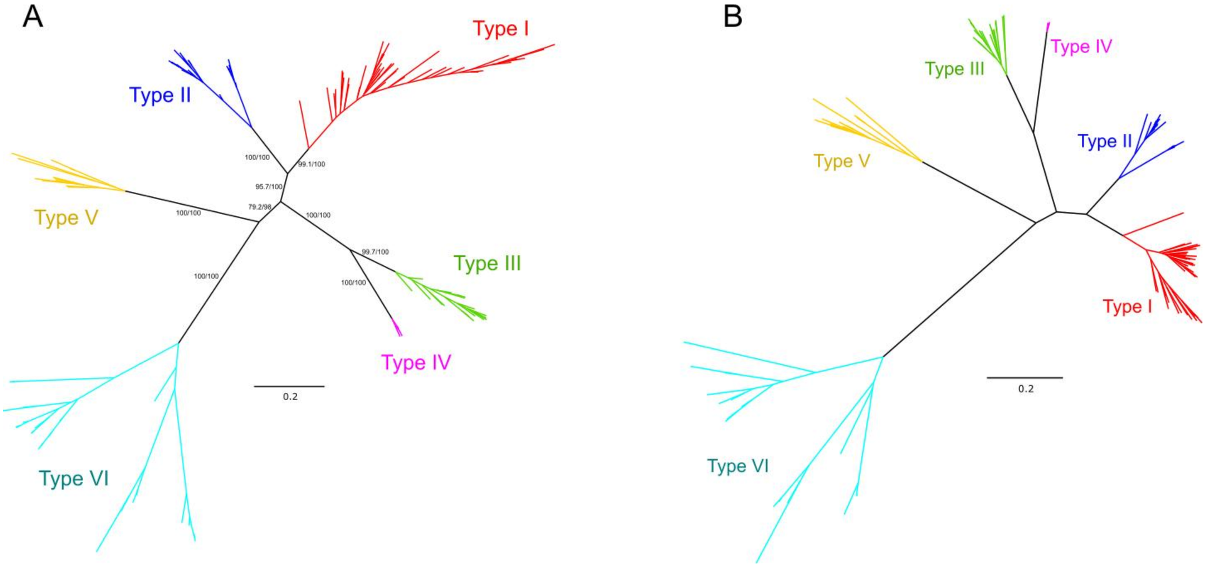


**Figure S5:** (A) Structure-informed maximum likelihood phylogeny of 166 Wmk homologs derived from 106 *Wolbachia* genomes. The tree was reconstructed using a partitioned framework in which protein sequence and structural alphabets were treated as separate partitions. The structural partition was modeled using 3DIPHY. Node supports (aLRT/BS) are based on 1000 replicates. Branches are colored according to the sequence-based Type classification shown in Fig. 2A and labeled accordingly.

(B) Phylogenetic tree inferred using FoldTree analysis of the same 166 structures. Branches are colored and labeled according to the sequence-based Type classification defined in Fig. 2A.

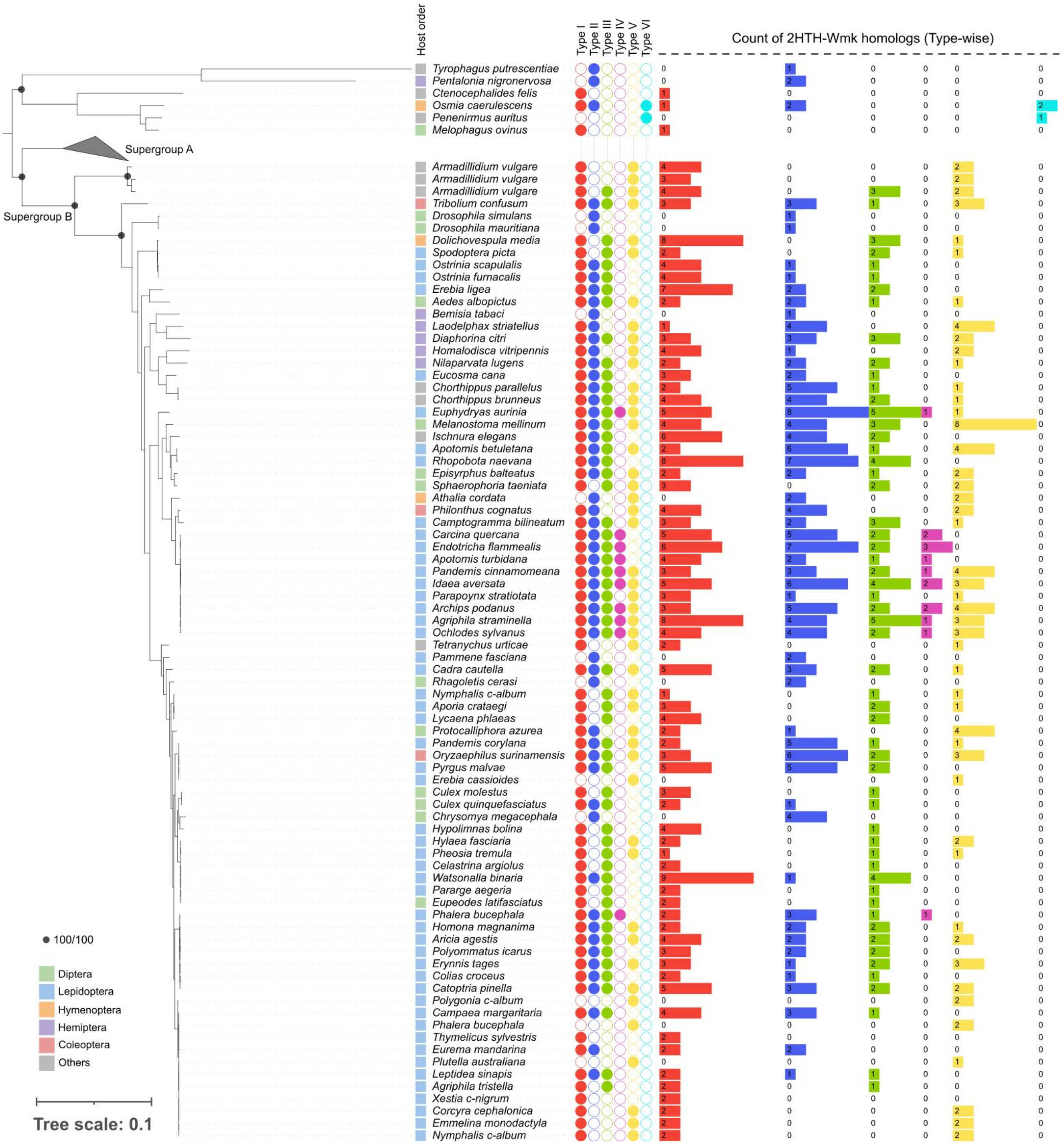


**Figure S6.** Pruned global ML phylogeny of *Wolbachia* retaining the 203 genomes that harbor at least one full-length Wmk homolog (Type I–VI). The original global phylogeny was reconstructed from 251 genomes. The tree here predominantly represents Supergroup B strains, along with members of Supergroups E and F. The clade corresponding to Supergroup A is collapsed here; an expanded view is provided in Fig. 3A. Node support values are shown as approximate likelihood ratio test and ultrafast bootstrap percentages (aLRT/BS) based on 1000 replicates, with support values displayed for deeper nodes only. Tip labels indicate host species name, host taxonomic order, and the presence and copy number of full-length Wmk homologs spanning Type I–VI.
